# Cancer cell elimination by cytotoxic T cell cooperation and additive damage

**DOI:** 10.1101/2020.04.22.054718

**Authors:** Bettina Weigelin, Annemieke Th. den Boer, Esther Wagena, Kelly Broen, Harry Dolstra, Rob J de Boer, Carl G. Figdor, Johannes Textor, Peter Friedl

**Affiliations:** Department of Cell Biology, RIMLS, Radboud University Medical Center, Nijmegen, The Netherlands; David H. Koch Center for Applied Research of Genitourinary Cancers, Department of Genitourinary Medical Oncology, The University of Texas MD Anderson Cancer Center, Houston, USA; Department of Preclinical Imaging and Radiopharmacy, Eberhard Karls University Tübingen, Germany; Department of Internal Medicine, Maastricht University, Maastricht, The Netherlands; Department of Laboratory Medicine - Laboratory of Hematology, Radboud University Medical Center, Nijmegen, The Netherlands; Theoretical Biology & Bioinformatics, Utrecht University, The Netherlands; Department of Tumor Immunology, RIMLS, Radboud University Medical Centre, Nijmegen, The Netherlands; Cancer Genomics Centre Netherlands (CGC.nl)

## Abstract

Cytotoxic T lymphocytes (CTL) eliminate tumor target cells in an antigen and cell-contact dependent manner. Lethal hit delivery occurs as a rapid and binary, “yes/no” process when immunogenicity is very high^1–3^, however *in vivo* CTL often fail to kill solid tumor cells during 1:1 conjugations^4–6^. Using long-term time-lapse microscopy in three distinct tumor cytotoxicity models and statistical modeling, we here show that migrating CTL transit between target cells and initiate apoptosis by a series of sublethal interactions (‘additive cytotoxicity’), while individual conjugations rarely induced apoptosis. Sublethal damage included perforin-dependent membrane pore formation, nuclear lamina rupture and DNA damage, and these events resolved within minutes to hours. In immunogenic B16F10 melanoma tumors *in vivo*, frequent serial engagements and sublethal hit delivery of CTL was largely confined to interstitial niches in the invasion front, resulting in eradication of invading tumor cells. Thus, additive cytotoxicity is a probabilistic process achieved by a series of CTL-target cell engagements and sublethal events. The need for additive “hits” has implications for the topographic mechanisms of elimination or immune evasion of tumor cells and microenvironmental regulation of CTL accumulation and cooperation by targeted therapy.

---

Cytotoxic T lymphocytes and NK cells can bind to and attack more than one target cell, termed “serial killing”^7–9^. Estimations from bulk killing assays and mathematical modeling suggest, that single CTL are capable of eliminating up to 20 target cells of the hematopoietic lineage per day both *in vitro* and *in vivo*^10,11^. This high efficacy of CTL-mediated serial killing, however, has not translated to the killing of solid tumors in mice^4,5,12^ and, likewise, is rarely observed in patients receiving adoptive transfer of tumor-specific TCR-engineered or CAR T cells^13^. To understand the discrepancy between CTL effector function against model target cell lines and solid tumors, we quantified the killing kinetics of single CTL in solid tumor models in 3D collagen-matrix based organotypic culture and the mouse dermis *in vivo*^14^. In these 3D environments, CTL mediated target cell killing depends on CTL motility and transient tumor interactions, not unlike kinetic priming models for CTL activation^15^.

## Serial conjugation and effector function

*In vitro* activated OVA-specific OT1 CTLs were confronted with transformed mouse embryonic fibroblasts expressing the OVA peptide (MEC-1/OVA) and the co-stimulatory molecule B7.1^13^. In contrast to tumor cells that evolved *in vivo*, this engineered model lacks natural immune escape modifications (e.g. down regulation of MHC-I or apoptosis resistance) and, thus, represents an idealized model for maximized CTL efficacy at a single cell level. After 30 h of co-culture, killing efficacy was near 100% at high effector-target (ET) ratios and reached background level below ET ratios of 1:128 (**Extended Data Fig. 1a**). To assess the serial killing efficacy of individual CTL, we analyzed antigen-specific CTL-target cell interactions and outcome in long-term time-lapse microscopy recordings by tracing individual CTL during interaction with target cells. The duration of individual CTL-target cell interactions was variable (lasting minutes to hours), with lag times between initial CTL binding and target cell death lasting 1.8 ± 1.5 h, and was followed by a subsequent period of ongoing CTL engagement with the dead cell body (‘necrophilic phase’) (**Extended Data Fig. 1 b-d**). This extended lag phase until apoptosis differs from lag phases obtained for CTL-mediated killing of target cells from leukocyte lineages, lasting <5 to 25 min^1–3^. In regions of low local CTL density, repetitive contacts resulted in the serial killing of multiple neighboring target cells by a single CTL (**Fig. 1a; Movie 1**). On the population level, 50% of the CTL acted as serial killers (maximum of 11 killed target cells/24 h), whereas a small CTL subset (15%) repeatedly contacted target cells without inducing apoptosis (**Extended Data Fig. 1e**). The percentage of CTL with killing capacity correlated with the surface expression of Lamp-1 by 85% of CTL, indicating recognition of the target cells and lytic vesicle exocytosis by the majority of CTL (**Fig. 1b**). The lag phase to apoptosis was neither compromised nor accelerated over consecutive killing events (**Fig. 1c**), which resulted in a consistent eradication frequency of 1 kill every 2 hours (**Extended Data Fig. 1f**). This excludes gain of cytotoxicity by kinetic priming through repetitive antigenic interactions. Thus, OT1 CTL serially eliminate highly immunogenic target cells over 24 h and in a non-exhaustive manner.

**Figure 1.**
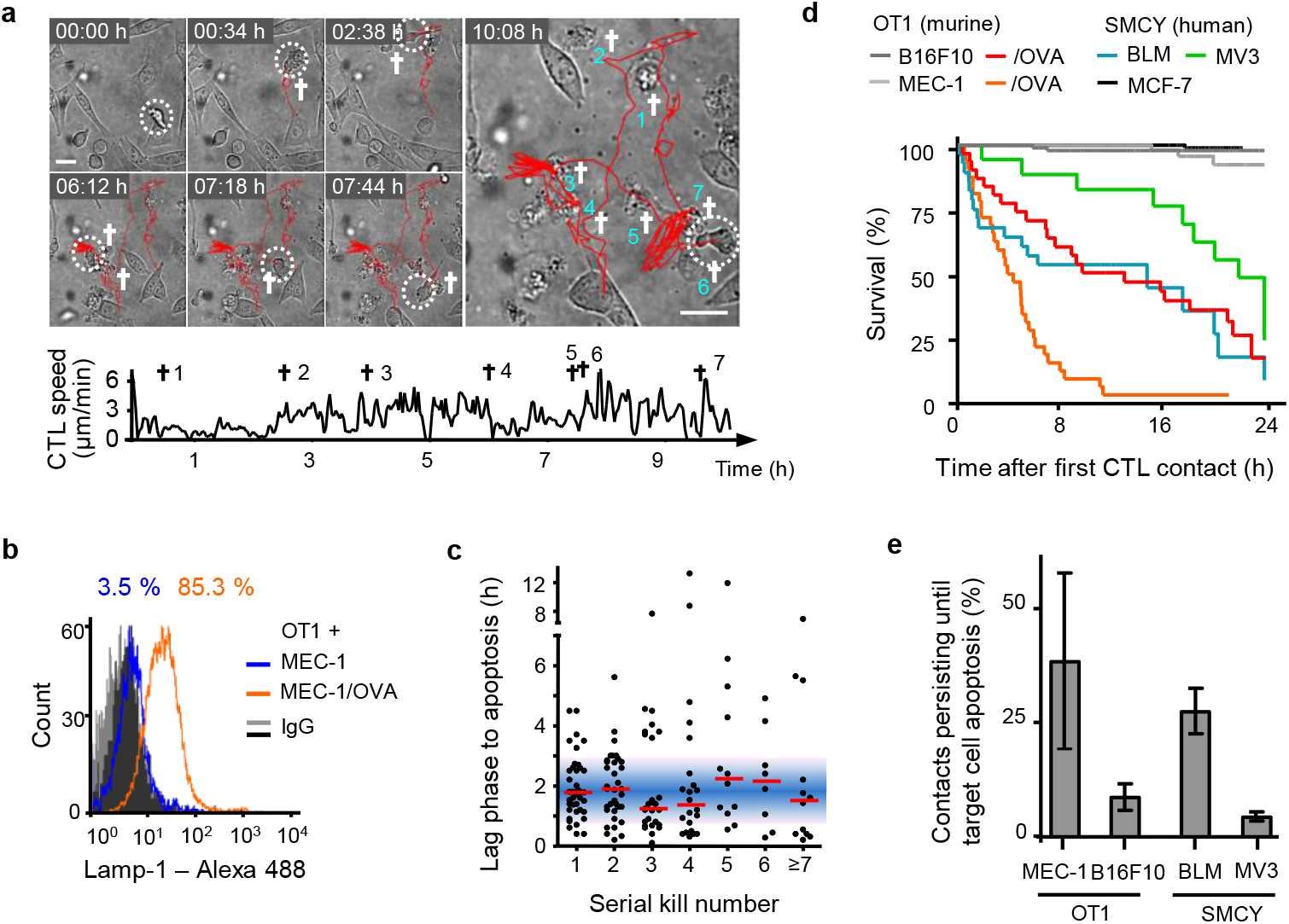
CTL serial conjugation and effector function. **a,** Time-lapse sequence and migration track of one OT1 CTL killing 7 MEC-1/OVA target cells sequentially within 11 h. Circles, CTL; cross, apoptotic target cell; scale bar, 20μm. **b,** Lamp-1 expression at the surface of OT1 CTL after 24-h of 3D coculture with MEC-1/OVA cells. Representative example of 3 independent experiments. **c,** Lag-phase until apoptosis of consecutive kills by the same CTL (43 CTL from 8 independent experiments). Red bars, median. **d,** Population survival of 4 distinct antigen-dependent target and 3 control models. Quantifications from 3 independent experiments for each cell line. **e,** Inefficiency of inducing apoptosis by individual CTL contacts in OT-1 and SMCY.A2 CTL models. Error bars, mean ± SD (≥ 100 contacts from ≥ 3 independent experiments per cell model).

## CTL induce sublethal damage

To compare effector function against solid tumor cells, which typically retain resistance to CTL mediated killing, OT1 CTL were confronted with mouse melanoma B16F10 cells expressing the OVA peptide (B16F10/OVA) (**Extended Data Fig. 2 a-d; Movie 2**). As a second model, IL-2 activated human SMCY.A2 CTL^14^, which recognize an HLA-A2 restricted antigen encoded on the y chromosome, were confronted with male human melanoma cell lines BLM or MV3 (**Extended Data Fig. 2 e-i; Movie 3**). Compared to the MEC-1/OVA cells, these three melanoma models show delayed, but ultimately effective target cell elimination at the end-point after 24 hours, whereas OVA-negative B16F10 or female MCF-7 cells survived (**Fig. 1e**). Thus, the endpoints of both murine and human models for probing CTL effector function show comparable target cell elimination in 3D culture. Notably, across all cell types tested, only a minority of individual CTL-target cell contacts induced apoptosis at first encounter, while 60-70% (MEC-1/OVA, BLM) or >90% (B16F10/OVA, MV3) of individual conjugations were followed by target cell survival (**Fig. 1e**).

CTL degranulation induces transient perforin-mediated pores in the target cell membrane, which facilitates diffusion of extracellular factors into the target cell, including CTL-derived granzyme B^16^ and extracellular calcium^17^. OT1 CTL deficient in perforin expression failed to kill B16F10/OVA cells in organotypic culture (**Extended Data Fig. 2j**), while CTL effector function remained intact after interference with Fas-FasL interaction (**Extended Data Fig. 2k**). Further, adding perforin-deficient OT1 CTL to a fixed number of wt OT1 CTL did not increase killing efficiency, indicating that perforin-independent mechanisms delivered by excess CTL, including soluble mediators, do not induce or enhance cytotoxicity against B16F10/OVA cells (**Extended Data Fig. 2l**). Thus, elimination of B16F10/OVA cells by OT1 CTL in 3D culture critically depends on perforin.

To visualize perforin-pore formation and to discriminate sublethal cytotoxic hits from functionally inert interactions, MEC-1/OVA and B16F10/OVA cells were engineered to express the calcium sensor GCaMP6s^18^, and monitored for transient Ca^2+^ influx through perforin pores (**Fig. 2a, panel 1; Extended Data Fig. 3a; Movie 4**). A substantial fraction of antigen-specific but non-lethal CTL-target cell contacts showed CTL associated intracellular Ca^2+^ events in B16F10/OVA cells (40%; **Fig. 2b, panel 2**), which were transient (median: 30 s; **Fig. 2c**) and in 90% followed by target cell survival (**Fig. 2d**). Ca^2+^ events originated at CTL - tumor cell contact regions, and differed from unspecific intracellular Ca^2+^ fluctuations by signal intensity and duration (**Extended Data Fig. 3 b,c**).

**Figure 2.**
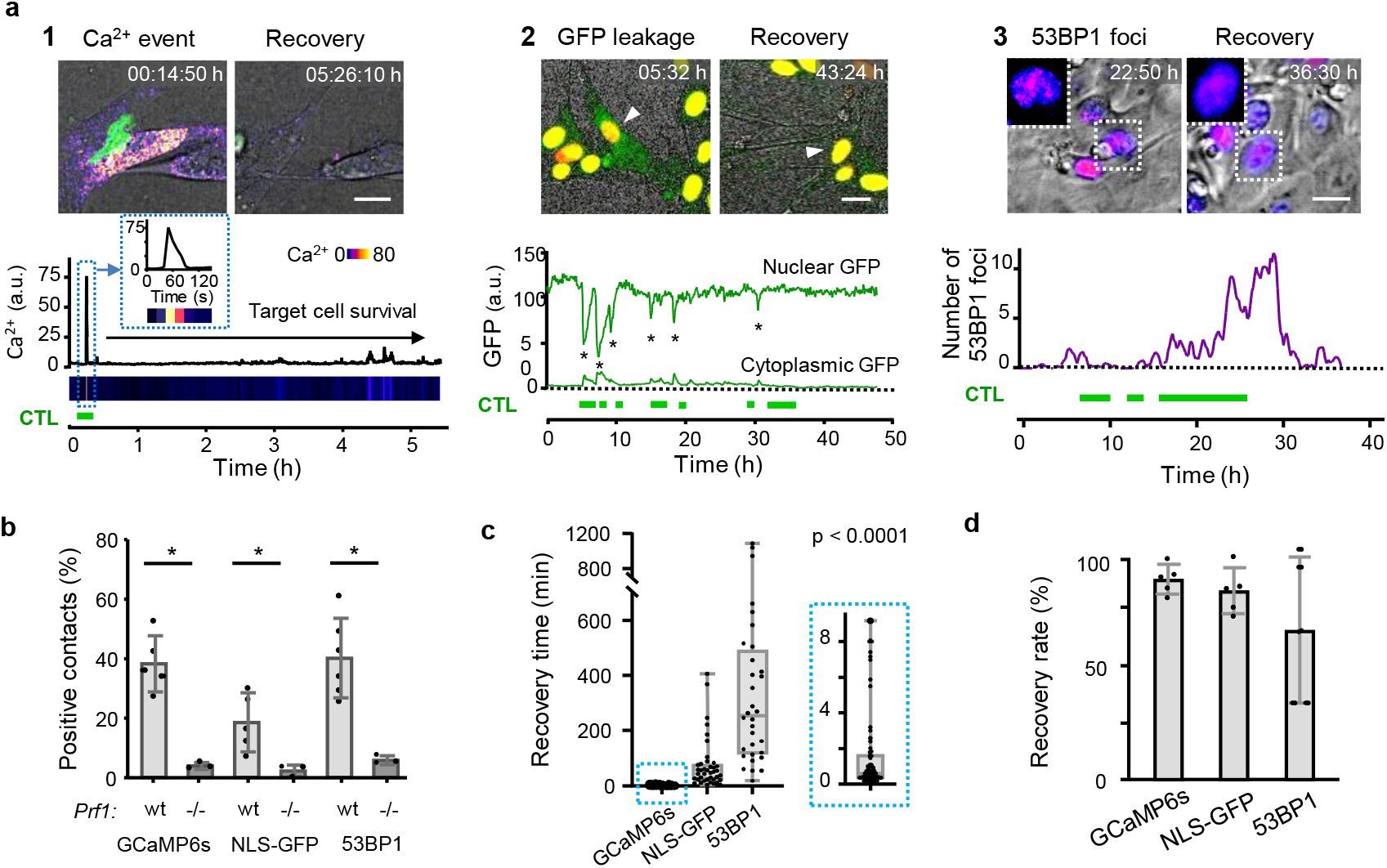
Sublethal damage induced by CTL. **a,** Reporter strategies. Green horizontal bars, duration of CTL contacts. Panel 1: CTL-mediated perforin pores visualized as Ca^2+^ influx into B16F10/OVA target cells using the GCaMP6s reporter. Time-resolved intensity plot of GCaMP6s event in the target cell cytosol followed by survival. Green, OT1 CTL (GFP); Fire LUT, Ca^2+^ level (GCaMP6s). Panel 2: CTL-mediated structural damage of the nuclear lamina monitored as NLS-GFP leakage into the cytosol. Nuclear and cytosolic GFP intensity plotted over time. Green, NLS-GFP; Red: Histone-2B-mCherry; Arrowhead: monitored cell. Asterisks, nuclear leakage events. Panel 3: CTL-mediated DNA damage response plotted as 53BP1trunc-Apple focalization in the nucleus over time. Fire LUT, 53BP1trunc-Apple. Insets, zooms of reversibility of 53BP1 foci. Bars, 10 μm. **b,** Percentage of contacts of wt and perforin-deficient CTL with B16F10/OVA cells inducing Ca^2+^ events, NLS-GFP cytosolic leakage and 53BP1trunc-Apple foci in target cells. Data show the mean ± SD from independent experiments: 3 (GCaMP6s /wt, NLS-GFP /wt /prf1−/−), 5 (53BP1trunc-Apple/wt) and triplicate movies from 1 experiment (GCaMP6s /prf1−/−, NLS-GFP /prf1−/−). *, p value < 0.05, two-tailed Mann-Whitney test comparing wt and prf1−/− datasets. **c,** Recovery times from initiation to termination of GCaMP6s, NLS-GFP and 53BP1trunc-Apple reporter activity. Box (25/75 percentile) and whiskers (minimum/maximum) of ≥ 32 cells (individual data points) from 3 (NLS-GFP, GCaMP6s) and 5 (53BP1trunc-Apple) independent experiments. p value, Kruskal-Wallis test comparing all groups corrected by Dunn’s multiple comparisons test. **d,** Percentage of cells with sublethal damage event which fully resolved. Data show the mean ± SD (5 independent experiments per reporter).

To test whether the variability of perforin release and consecutive Ca^2+^ events in B16F10/OVA cells were a consequence of heterogeneous TCR engagement, we quantified Ca^2+^ signals in OT1 CTL upon target cell contact. OT1 CTL showed comparably high rates of Ca^2+^ signaling when contacting MEC-1/OVA and B16F10/OVA cells (80% to 85%), typically within seconds after contact initiation (**Extended Data Fig. 3 d**). When co-registered with Ca^2+^ events in target cells, 40% of Ca^2+^ positive-CTL contacts with B16F10/OVA coincided with, or were immediately followed by a Ca^2+^ event in the target cell (**Extended Data Fig. 3 e,f**). In conclusion, while TCR triggering in OT1 CTL occurs reliably, the induction of perforin-events in the target cell varied.

## Structural damage in target cells

To address whether transient transmembrane pores were associated with structural intracellular damage, B16F10/OVA cells were engineered to express NLS-GFP^19^ or 53BP1trunc-Apple^20^. NLS-GFP leakage into the cytoplasm was detected in 25% of CTL-target cell contacts and absent when CTL lacked perforin expression **(Fig. 2a, panel 2; b, Extended Data Fig. 3 g-j; Movie 5)**. Recovery, as indicated by diminishing NLS-GFP signal in the cytosol and recovery in the nucleus, occurred in 75% of events within minutes to hours (median: 49 min; **Fig. 2 c,d**). 53BP1 initiates DNA damage repair complexes, which can be visualized as repair foci by 53BP1trunc-Apple^20^ (**Extended Fig. 3 k; Movie 6)**. 53BP1 foci were induced in 35% of CTL contacts, in dependence of perforin expression in OT1 CTL (**Fig. 2 a,b**) and CTL density (**Extended Fig. 3 l,m**). CTL-induced 53BP1 foci persisted for several hours (median: 4 h) and were resolved in 73% of events (**Fig. 2 c,d**). These data indicate that CTL contacts induce reversible sublethal damage to the nuclear lamina and DNA.

## Death induction by multiple CTL

Across all tested tumor models, sequential or simultaneous contacts by multiple CTL with the same target cell occurred before target cell death (**Fig. 3a**). In the B16F10 model, >90% of successful kills were preceded by 2-9 CTL encounters by distinct CTL (**Extended Data Fig. 4a**). When Ca^2+^ events in target cells were recorded, death induction was preceded by serial Ca^2+^ events with variable onset and frequency between hits (**Fig. 3b; Movie 4, 7**). The lag time between contact initiation and first Ca^2+^ event was 6 min in MEC-1/OVA cells and 16 min in B16F10/OVA cells (**Extended Data Fig. 4b**). Killing of the MEC-1/OVA cells was preceded by Ca^2+^ events with 5 min median interval, while Ca^2+^ events in B16F10/OVA cells occurred with longer intervals of 20 min (**Extended Data Fig. 4c**). While >50% of contacts induced only one Ca^2+^ event in B16F10/OVA cells, in 40% of CTL contacts yielded at least 2 repetitive Ca^2+^ events (**Extended Data Fig. 4d)**. Thus, compared to MEC-1/OVA cells, delayed killing of B16F10/OVA cells correlated with delayed delivery of Ca^2+^ events (**Extended Data Fig. 4e)**.

**Figure 3.**
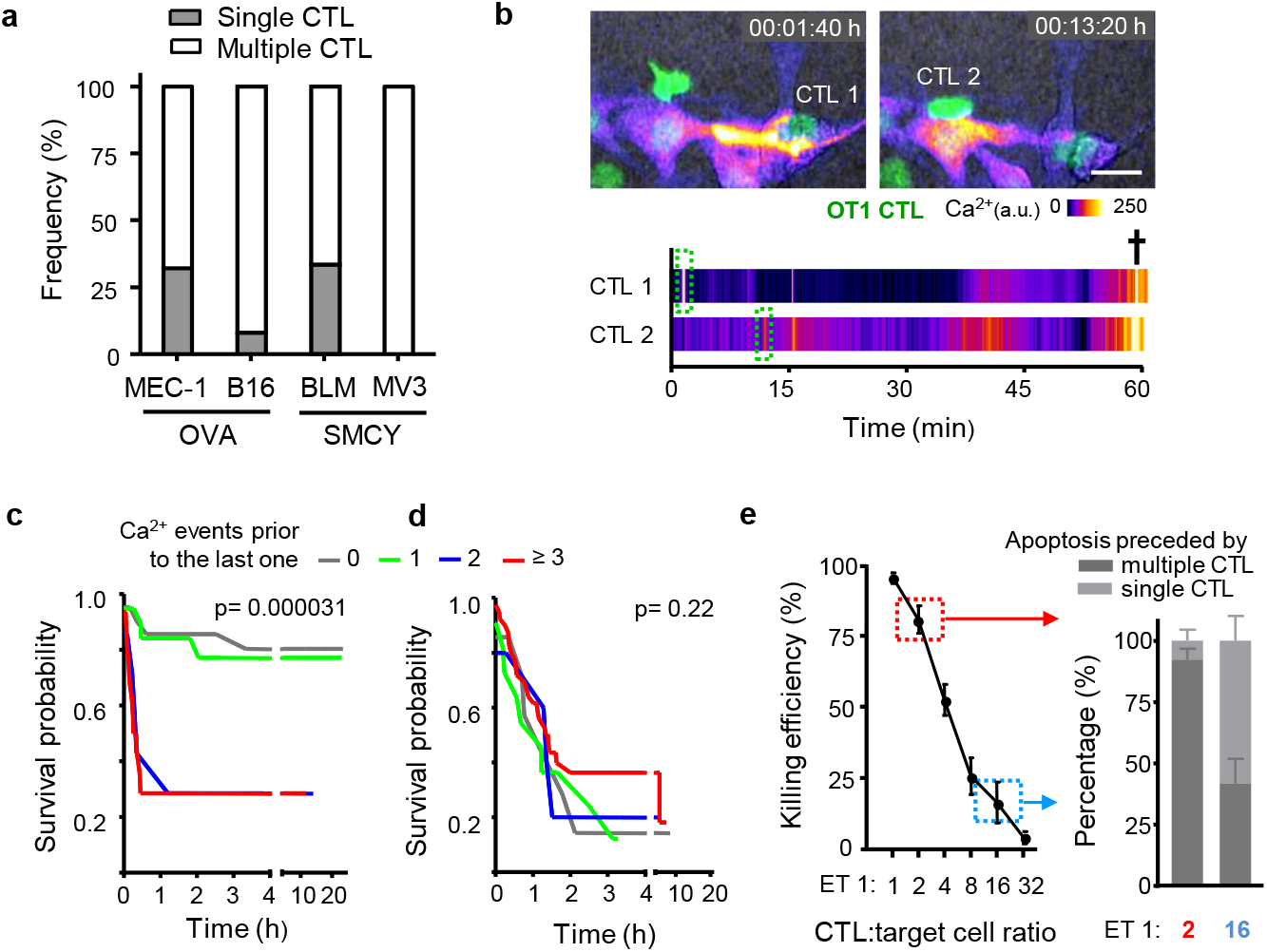
Additive cytotoxicity. **a,** Percentage of multiple contacts preceding apoptosis in murine and human melanoma models. Pooled data representing ≥100 contacts from ≥3 independent experiments per cell line. **b,** Time-lapse sequence and intensity plot of multiple Ca^2+^ events followed by target cell apoptosis. Green fluorescence, OT1 CTL (dsRed); Fire LUT, Ca^2+^ intensity (GCaMP6s). Cross, target cell death; Scale bar, 20 μm. **c,** Survival probability of B16F10/OVA cells having received increasing numbers of Ca^2+^ events. **d,** Simulation of stochastic apoptosis induction by permutation of waiting times between Ca^2+^ events, survival and lag times until apoptosis. p-values in (c, d), log rank test comparing all groups. **e,** Killing efficacy of B16F10/OVA cells and percentage of preceding single or multiple interacting CTL in dependence of ET ratio. Left panel, means and SD from 3 independent experiments; right panel, 110 contacts from 4 independent experiments per ET ratio.

## Additive cytotoxicity

We then addressed whether apoptosis could have been induced by rare deadly CTL hits (‘stochastic killing’) instead of sublethal contacts that add up over time (‘additive cytotoxicity’). Therefore, we analyzed whether the lethality of the final hit was enhanced by, or independent of, previous Ca^2+^ events, by plotting the lag time to apoptosis and target cell survival probability in relation to the number of pre-final Ca^2+^ events. Target cells which received two or more hits prior to the lethal one showed accelerated apoptosis induction, together with a sharp decrease in survival probability (**Fig. 3c; Extended Data Fig. 5a,b**). The dependence of the lag time to apoptosis on prior CTL hits is expected when additive effects exists, but it is inconsistent with the stochastic killing hypothesis. To explore stochastic killing directly, we performed the same analysis on simulated data, using randomly swapped times between hits and between the last hit and apoptosis. Here, cell death induction was gradual, and neither the lag time to apoptosis nor the survival probability was dependent on the number of prior hits (**Fig. 3d**).

To address how long sublethal events remain relevant, we estimated the time required to repair the damage caused by a single Ca^2+^ hit, using a Cox regression model based on additive killing. This resulted in an estimated damage half-life of 56.7 minutes (95% CI: 33.1-112.2) of Ca^2+^ events to contribute to lethal outcome. This interval was consistent with the recovery kinetics of nuclear lamina defects (**Fig. 2c**). Using the Bayesian Information Criterion (BIC) to compare model fits, showed that the model which integrates serial damage and decay explained the data better (BIC difference >10) than a model based on the number of Ca^2+^ events alone.

## Multi-hit delivery as a function of CTL density

To test whether apoptosis induction under conditions of high CTL density facilitated additive, multi-hit interactions or, instead, a higher probability of lethal single-hit interactions of few CTL^21^, we titrated CTL density and monitored individual CTL-target cell interactions by time-lapse microscopy. At high CTL density (ET 1:2) target cell death was frequent and preceded by multiple CTL interactions (**Fig. 3e**), whereas at low CTL density (ET 1:16) infrequent apoptotic events were near-exclusively preceded by single-contact engagements (**Fig. 3e**). The CTL density effect was not enhanced by the addition of perforin-deficient CTL (**Extended Data Fig. 2l**). This indicates that high CTL density (“swarming”) enables efficient apoptosis induction by favoring serial perforin-dependent hits, whereas the poor killing at low CTL density largely relies upon single encounters.

## Sublethal hit delivery *in vivo*

To address whether multiple encounters by CTL mediate anti-tumor cytotoxicity *in vivo*, activated OT1 CTL were adoptively transferred into C57BL/6 J mice bearing intradermal B16F10 melanoma and monitored longitudinally by intravital microscopy through a skin window (**Extended Data Fig. 6 a, b**). A single application of OT1 CTL caused a transient, dose-dependent growth delay of the OVA antigen-expressing but not of control tumors (**Extended Data Fig. 6c).** B16F10 tumors invade the dermis as multi-cellular strands^22^. Concomitantly, OT1 CTL first accumulated in the tumor periphery and subsequently redistributed towards the invasive tumor front (**Fig. 4a**). This resulted in local ET ratios of 1:1 along the tumor-stroma interface, whereas ET ratios in the tumor core remained below 1:250 (**Extended Data Fig. 6 d,e)**. To identify where and by which contact mechanism eradication of tumor cells was achieved, CTL contacts and outcome were mapped using histone-2B/mCherry (H2B/mCherry) to detect nuclear fragmentation in B16F10/OVA cells^23^. Despite comparable ET ratios, high fragmentation rates occurred in the invasion niche but not the tumor rim (**Fig. 4b**). In either zone, >95% of CTL contacts were transient, short-lived (median duration 15 min) and non-lethal (**Fig. 4c**). When aggregated, >86% of apoptotic events were preceded by multiple CTL contacts, and a minority (<14%) by individual CTL conjugation (**Fig. 4d**). In the tumor invasion niche, high local CTL density coincided with confined migration along tissue structures with enhanced speed compared to the main tumor mass (**Extended Data Fig. 7a, b; Movie 8**). This supported frequent contacts to B16F10/OVA cells with cumulative contact duration reaching >1h and nuclear fragmentation in tumor cells in a time-dependent manner (**Fig. 4e, Movie 9**).

**Figure 4.**
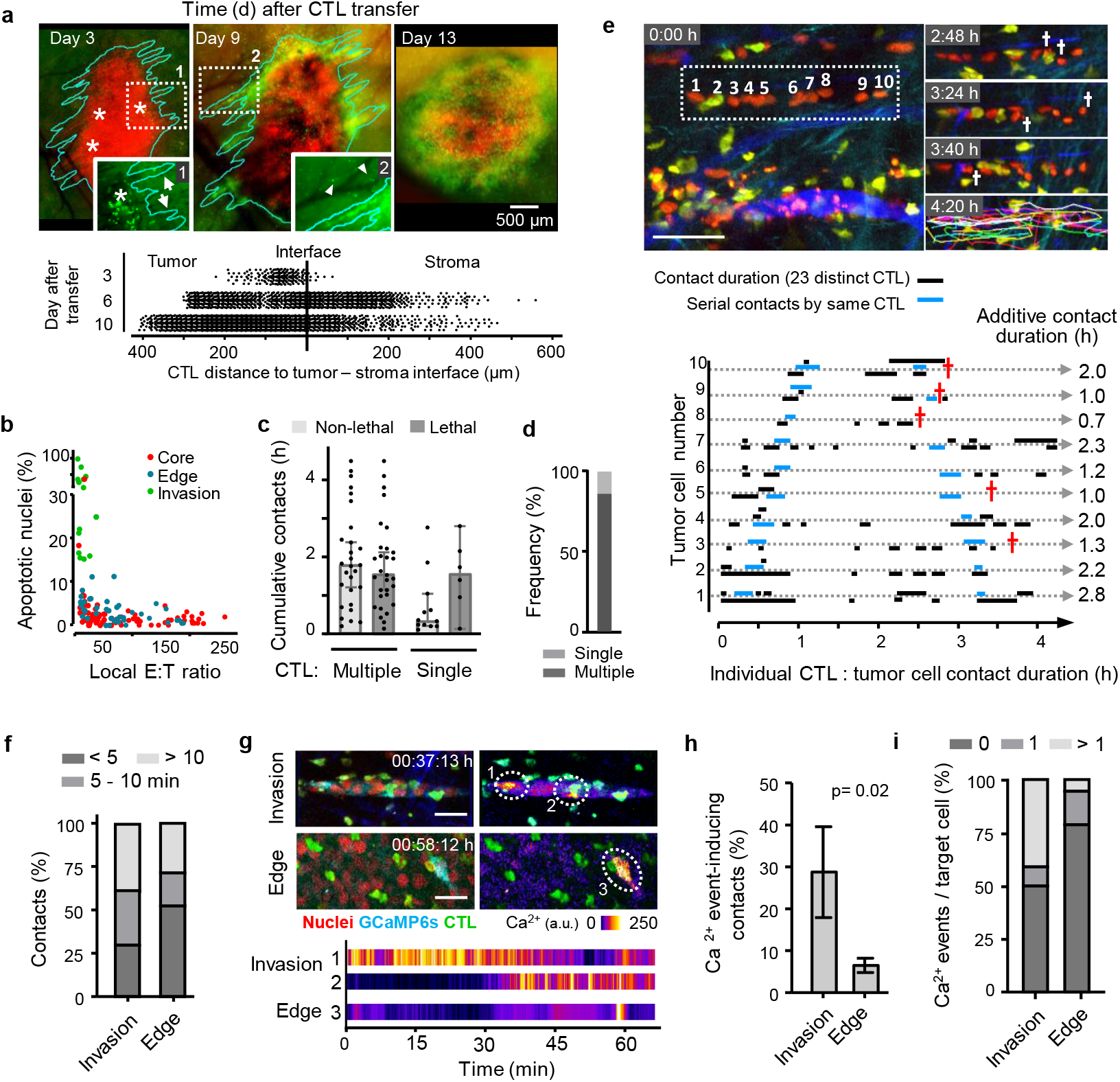
Additive cytotoxicity in live tumors. **a,** Time-dependent CTL accumulation along the tumor-stroma interface. Red, nuclei B16F10/OVA cells; green: OT1 CTL. Lower panel, position of individual CTL. Representative example of one tumor. **b,** Correlation of CTL density and apoptotic frequency in tumor subregions. Pooled data, 8 independent mice. **c,** Cumulative contact duration and outcome of single or multiple CTL contacting B16F10/OVA cells. **d,** Frequency of apoptosis induction by single or multiple CTL. Data in (c, d) represent 40 non-lethal and 37 apoptotic events from 150 h of movies pooled from 20 independent tumors. **e,** Representative micrographs and quantification of serial engagements of multiple CTL with B16F10/OVA cells and outcome. Red cross, target cell apoptosis. 10 target cells from one tumor. **f,** Percentage of contacts between CTL with B16F10/OVA cells and duration category in the invasion zone versus tumor edge. **g,** Images from time-lapse recordings of GCaMP6s activity in B16F10/OVA target cells in distinct tumor subregions. Dotted circles indicate the example areas plotted for GCaMP6s intensity in the graph below. **h,** Percentage of CTL contacts which induced one or more Ca^2+^ events in invasion zones versus CTL-rich subregions at the tumor edge. **i,** Percentage of tumor cells receiving none, one or more than one Ca^2+^ event within a cumulative observation time of 3 h per tumor subregion. Data in (f, h, i) represent 228 contacts from 4 independent mice. Data in (h), mean ± SD (3 positions per subregion from 4 independent mice). Scale bars (e,g), 50 μm.

To discriminate functional from non-functional, irrelevant interactions, we analyzed the occurrence of Ca^2+^ events using GCaMP6s expressing tumors. Most Ca^2+^ events (80%) in B16F10/OVA cells were associated with CTL contacts, but rare without interacting CTL (**Extended Data Fig. 7 c,d**). In invasive tumor subregions with high apoptosis rates, the frequency of CTL contacts inducing Ca^2+^ events was 5-fold increased, compared to the non-invading tumor rim (**Fig. 4 g,h; Movie 10, 11**). Thus, *in vivo* eradication of tumor regions by CTL is a function of local CTL density and frequent sublethal interactions.

## Conclusions

Compared with the deterministic and reliable elimination of B cells^1–3^, CTL effector function against solid tumor cells is inefficient, with a high failure rate, and rarely completed by a single CTL but dependent on a sequence of sublethal hits. Consequently, apoptosis induction is not a binary event but instead implies the probabilistic accumulation of a death signal within the target cell over time, not unlike the accumulation of activation signal in naïve T cells by successive encounters with APCs^24–26^. The reversible structural damage in the plasma and nuclear membrane and DNA integrity induced by CTL resembles damages induced by chemical and mechanical assault^19,27,28^ which are also incremental with challenge and reversible by repair. Thus, irrelevant interactions may alternate with variably damaging events and become integrated over time in the target cell until apoptosis is induced or recovery achieved (**Extended Data Fig. 9**). Additive cytotoxicity may provide a mechanism by which dense CTL infiltrates induce apoptosis in a viral infection model^29^ or, here, the tumor invasion zone. Alternatively, inefficient sublethal hit delivery may explain failed eradication despite high CTL numbers in solid tumors in mice^5^, during alloimmune response against transplants^30^ or, here, subregions of the tumor margin.

CTL density control may, thus, provide a filter limiting accidental cell damage by single, miss-targeted CTL. Additive cytotoxicity may enable cooperation of CTL with differing TCR-specificity or different cytotoxic recognition strategies, such as CTL and NK cells. Lethal hit delivery by serial CTL engagements, as gradual, tunable process, may further respond to microenvironmental and therapeutic immunomodulation, including stabilization of CTL-target cell contacts^31,32^, modulating local concentration of soluble factors^33^, activation of immunostimulatory pathways^34^ and/or blockade of immune checkpoints^35^. In conclusion, serial conjugation and delivery of sublethal hits define the efficacy of CTL effector function which can be exploited by targeted therapy to increase both single contact efficacy and CTL cooperativity.

## Methods summary

### Organotypic three-dimensional cytotoxicity assay and time lapse microscopy

Sub-confluent target cell monolayers were overlaid with 3D fibrillar collagen (PureCol, concentration: 1.7 mg/ml) containing *in vitro* activated OT1 CTL (cell models and activation protocols described in the Methods section). CTL migration, interactions with target cells and apoptosis induction was monitored by time-lapse bright-field microscopy frame intervals of 30 sec for 24 h followed by manual cell tracking and quantitative population analysis.

### Monitoring sublethal damage

Target cells were lentivirally transduced to stably express the calcium sensor GCaMP6^17^, NLS-GFP^19^ or 53BP1trunc-Apple^36^. OT1 CTL-target cell conjugation and consecutive reporter activity were coregistered by confocal 3D time-lapse microscopy (Leica SP8 SMD) at 488 nm and 561 nm excitation (both 0.05 mW) at frame intervals ranging from seconds to minutes and total observation periods of up to 30 h (GCaMP6s: 10 sec/12 h; NLS-GFP: 2 min/30 h; 53BP1: 5 min/40 h). Intravital microscopy of GCaMP6s activity was performed by simultaneous scanning with 910 nm (eGFP and Alexa750, 15 mW) and 1140 nm (mCherry, dsRed and SHG, 30 mW) with a sampling rate of 1 frame / 10-15 sec over periods of 1-2 h. CTL position and reporter activity were obtained by manual or semi-automated image segmentation and quantification using ImageJ/FIJI.

### Intravital multiphoton microscopy

Histone-2B/mCherry expressing B16F10/OVA melanoma cells (1×10^5^) were injected into the deep dermis of C57/B16 J mice (Charles River) carrying a dorsal skin-fold chamber and were repeatedly monitored for up to 15 days^28^. With the onset of tumor invasion and angiogenesis at day 3 after implantation, *in vitro* activated eGFP or dsRed OT1 CTLs (0.5-1 ×10^6^) were injected intravenously. Multi-parameter intravital multiphoton microscopy was performed on anesthesized mice (1-3% isoflurane in oxygen) on a temperature-controlled stage (37°C) recording up to 5 channels simultaneously to visualize blood vasculature (70kD dextran/ AlexaFluor750) and peri-tumor stroma (SHG).

### Statistical modeling

Survival curves of target cells receiving serial Ca^2+^ events were computed in GNU R using the ‘survival’ and ‘rms’ packages.

### Statistical analysis

Unpaired student’s t-tests or Mann-Whitney U-tests, as appropriate, were applied using GraphPad Prism 8.

## Supporting information

Supplementary Method Section

**Supplementary Information** is linked to the online version of the paper at www.nature.com/nature.

## Acknowledgements

This work was supported by the Dutch Cancer Foundation (KWF 2008-4031) to C.G.F.. and P.F., NWO-Rubicon (019.162LW.020) to BW, a personal KWF grant to AdB), the FP7 of the European Union (ENCITE HEALTH TH-15-2008-208142), NWO-VICI (918.11.626), the European Research Council (617430-DEEPINSIGHT), and the Cancer Genomics Cancer, The Netherlands to P.F.. Time-lapse confocal microscopy was enabled by an NWO investment grant (834.13.003). We thank Stephen P. Schoenberger for providing the MEC-1/OVA cell line.

## Author contributions

B.W., P.F. designed the experiments, interpreted the data and wrote the paper. A.d.B. designed and performed experiments, B.W., E.W., quantitatively analyzed the data, K.B. isolated, characterized and cultured the human SMCY.A2 CTL, J.T. and R.d.B performed mathematical analyses, H.D. and C.F. contributed to data interpretation. All authors read and corrected the manuscript.

## Author Information

Reprints and permissions information is available at www.nature.com/reprints. The authors declare no competing financial interests. Correspondence and requests for materials should be addressed to.

## Supplementary videos

**Supplementary movie 1.** Serial killing of seven MEC-1/OVA target cells by one OT1 CTL (3D collagen assay). Time, hours:min. Field size 80 × 80 μm.

**Supplementary movie 2.** B16F10/OVA mouse melanoma target cells cocultured with OT1 CTL. Time, hours:min. Field size 340 × 280 μm.

**Supplementary movie 3.** BLM human melanoma target cells cocultured with SMCY.A2 CTL. Time, hours:min. Field size 280 × 270 μm.

**Supplementary movie 4.** B16F10/OVA target cell expressing the Ca^2+^ sensor GCaMP6s (Fire LUT) contacted by an OT1 CTL (green). A single short-lived Ca^2+^ event followed by target cell survival (part 1). Repetitive Ca^2+^ events preceding target cell apoptosis (part 2). Time, hours:min:sec. Field size 60 × 55 μm (part 1), 60 × 45 μm (part 2).

**Supplementary movie 5.** Confocal time sequence of B16F10/OVA target cell expressing the NLS-GFP reporter (green) and H2B-mCherry (red) during contact by OT1 CTL (unlabeled, brightfield channel), causing sequential NLS-GFP leakage events with or without apoptosis induction. Circles indicate leakage events. Time, hours:min. Field size 180 × 180 μm.

**Supplementary movie 6.** Confocal time sequence of B16F10/OVA target cell expressing the 53BP1trunc-Apple reporter (Fire LUT) attacked by OT1 CTL (unlabeled), causing 53BP1trunc-Apple focalization followed by resolution (part 1) or apoptosis induction (part 2). Arrowheads indicate DNA repair foci. Time, hours:min. Field size 60 × 60 μm (part 1), 80 × 80 μm (part 2).

**Supplementary movie 7.** Serial engagements of multiple OT1 CTL (green) with B16F10/OVA target cell expressing the Ca^2+^ sensor GCaMP6s (Fire LUT) followed by target cell apoptosis. Time, hours:min:sec. Field size 140 × 85 μm.

**Supplementary movie 8.** Dynamic sublethal conjugations of tissue invading B16F10/OVA cells expressing Histone-2b-mCherry (Red) by OT1 CTL (dsRed2, green). Perfused blood vessels (Al750, blue), Collagen fiber (SHG, red). hours:min. Field size 220 × 90 μm.

**Supplementary movie 9.** Serial engagements of multiple OT1 CTL (dsRed2, yellow) with invading B16F10/OVA target cells (H2B-mCherry nuclei, red) followed by apoptosis of several target cells after multiple CTL contacts. Perfused blood vessels (Al750, blue), Collagen fiber (SHG, cyan). Arrow heads indicate nuclear condensation as first sign of apoptosis. hours:min. Field size 220 × 170 μm.

**Supplementary movie 10.** B16F10/OVA cells expressing the Ca^2+^ sensor GCaMP6s (Cyan/ Fire LUT) in the tumor rim, infiltrated by OT1 CTL (dsRed2, green). CTL conjugation cause sublethal hits in few tumor cells. Perfused blood vessels (Al750, blue), Collagen fiber (SHG, red). Circles indicate Ca^2+^ events. hours:min:sec. Field size 330 × 330 μm.

**Supplementary movie 11.** Tissue invading B16F10/OVA cells expressing the Ca^2+^ sensor GCaMP6s (Cyan/ Fire LUT) serially contacted by OT1 CTL (dsRed2, green). Dynamic CTL conjugations cause repetitive Ca^2+^ events in a high percentage of tumor cells. Perfused blood vessels (Al750, blue), Collagen fiber (SHG, red). Circles indicate Ca^2+^ events. hours:min:sec. Field size 160 × 120 μm.

**Extended data figure 1.**
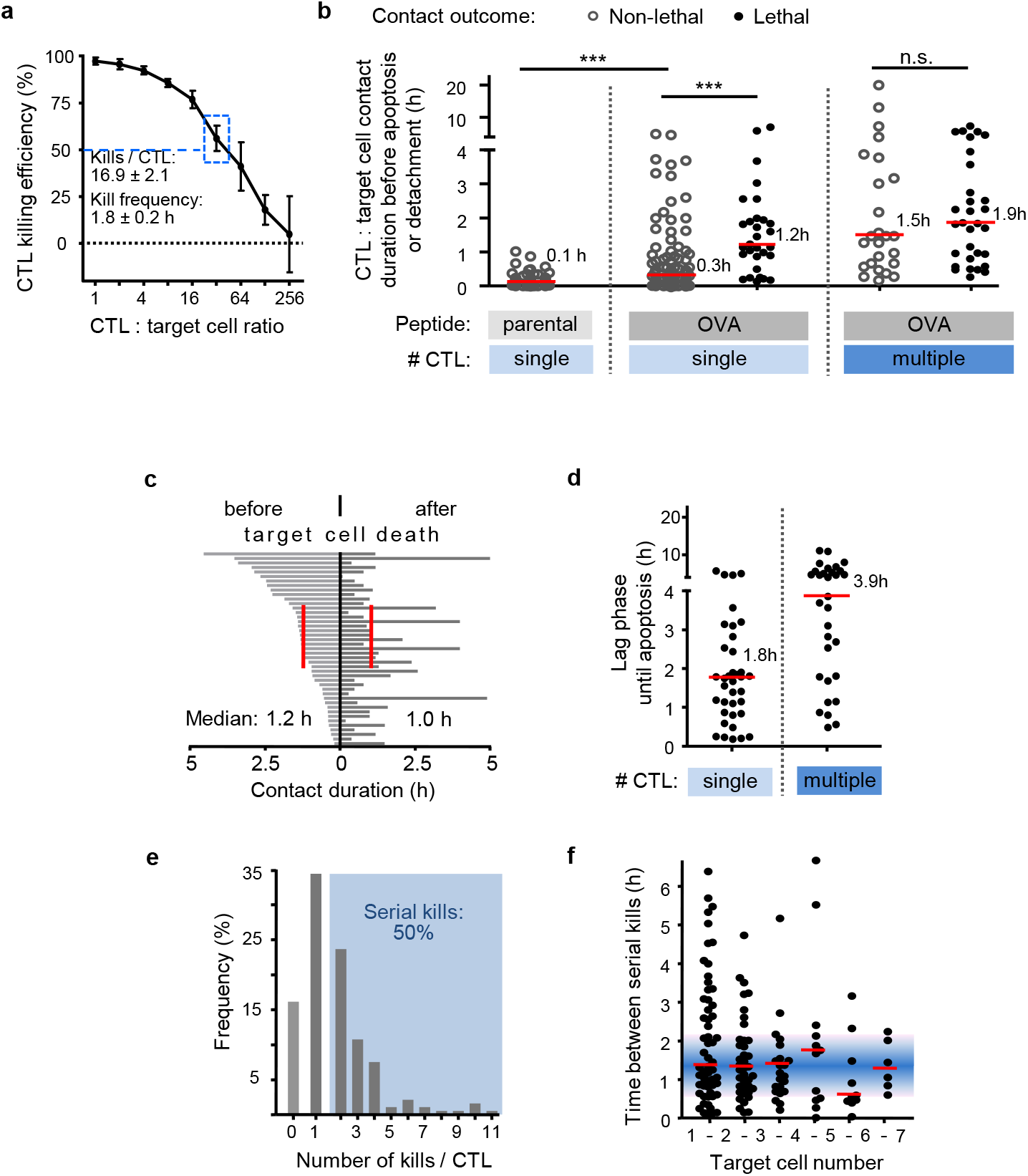
CTL contact phases, types and duration in MEC-1/OVA model. **a,** Killing of MEC-1/OVA targets by OT1 CTL at different ET ratios, determined by flow cytometry after 24 h of co-culture. Data show means ± SD (3 independent experiments). **b,** Cumulative contact duration between CTL andtarget cell and contact outcomes. **c,** Duration of CTL-target cell contacts before and after apoptosis. **d,** Lag phase until apoptosis of target cells induced by single or multiple CTL. **e,** Frequency of serial kills. **f,** Time between serial killing events mediated by the same CTL. *** p < 0.001; n.s., not significant (two-tailed Mann-Whitney test). Red bars, median. Data in (**b-f**) were pooled from 9 independent experiments.

**Extended data figure 2.**
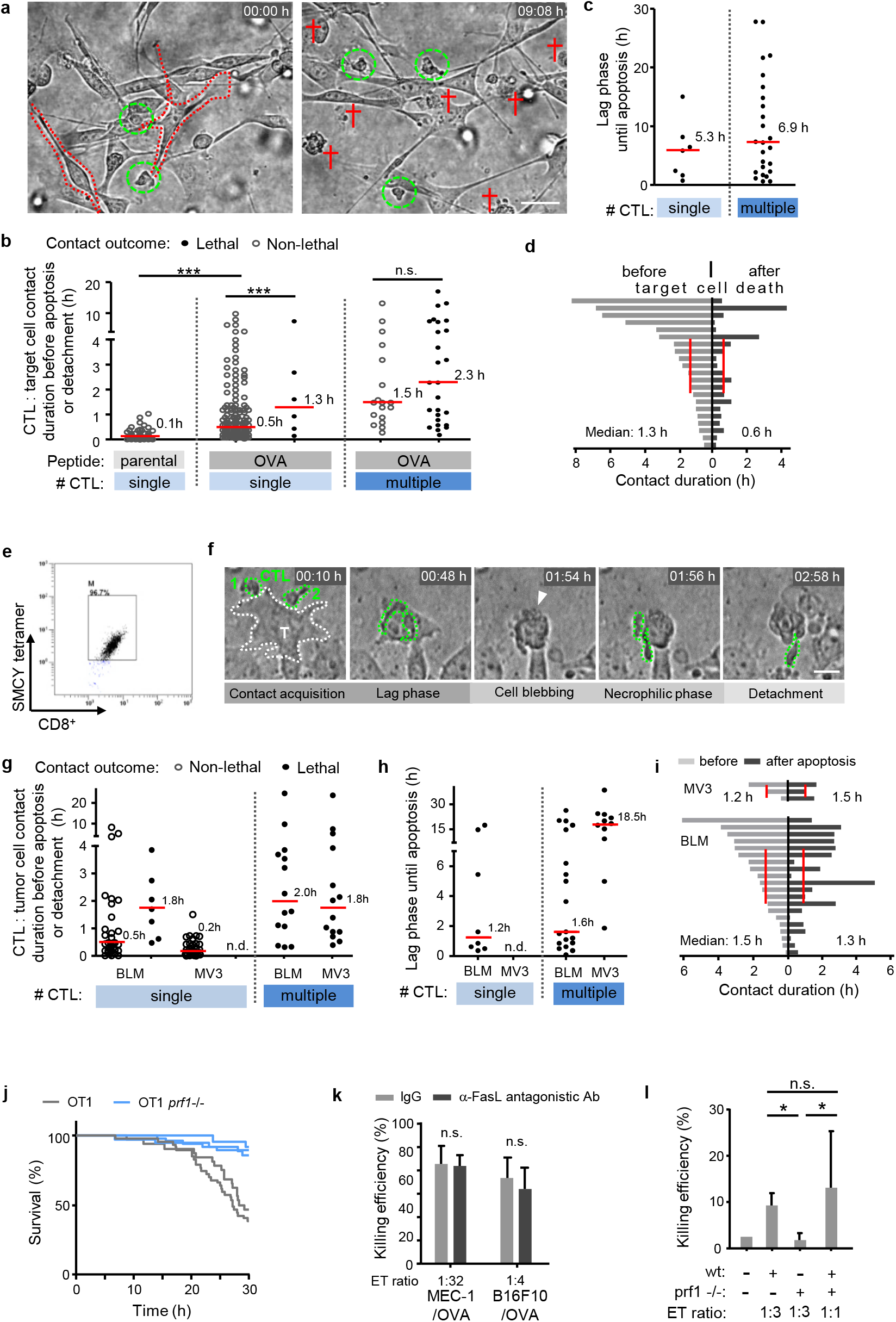
CTL effector function against mouse B16F10 and human MV3 and BLM melanoma cells. **a,** Overview image of sub-confluent monolayer of B16F10/OVA with CTL in the 3D collagen matrix interface assay at consecutive time points. Green circles, CTL; red cross, apoptotic target cell. Scale bar, 50 μm. **b,** Duration of contacts between individual and multiple CTL with B16F10/OVA target cells from initiation until apoptosis induction. **c,** Lag phase until apoptosis induced by contacts from single or multiple, sequentially engaged CTL. **d,** Duration of contact phases between individual CTL with target cell before and after target cell blebbing, obtained by manual tracking. Red lines, median. Data in (b-d) show 232 contacts pooled from 5 independent experiments. **e,** SMCY.A2 tetramer / CD8 double staining confirming CTL specificity and purity of > 95%. **f,** Time-lapse sequence of two simultaneously engaging CTL with human BLM melanoma target cell followed by target-cell apoptosis. Arrowhead, blebbing target cell during death. Scale bar, 20 μm. **g,** SMCY.A2 CTL contact durations with BLM and MV3 melanoma targets and contact outcome. n.d., not detected. **h,** Lag phase until apoptosis induced by single or multiple CTL. **i,** CTL contact duration with the same target cell before and after target cell blebbing. Red bars, median. Data in (g-i) show 50 (MV3) and 53 (BLM) contacts pooled from 3 (MV3) and 4 (BLM) independent experiments. **j,** CTL-mediated killing efficiency of B16F10/OVA cells by wild-type (wt) and perforin-deficient (prf1 −/−) OT1 CTL in 3D culture. Data acquisition and analyses as in Fig. 1d. Data show duplicates of one experiment. **k,** CTL killing efficiency upon blocking FasL by α-FasL Ab (MFL4) (10μg/ml). Data represent the means ± SD (n=3). n.s., non-significant (two-tailed Mann-Whitney test). **l,** B16F10/OVA Histone-2B-mCherry / NLS-GFP cocultures with wild type (wt) and mixed, wt and prf1 −/− OT1 CTL were monitored for 40 h by confocal time-lapse microscopy. Readout of CTL-mediated apoptosis induction was achieved by manual analysis of nuclear condensation and fragmentation. Data represent the means ± SD (3 independent experiments). *, p < 0.01.

**Extended Data Figure 3:**
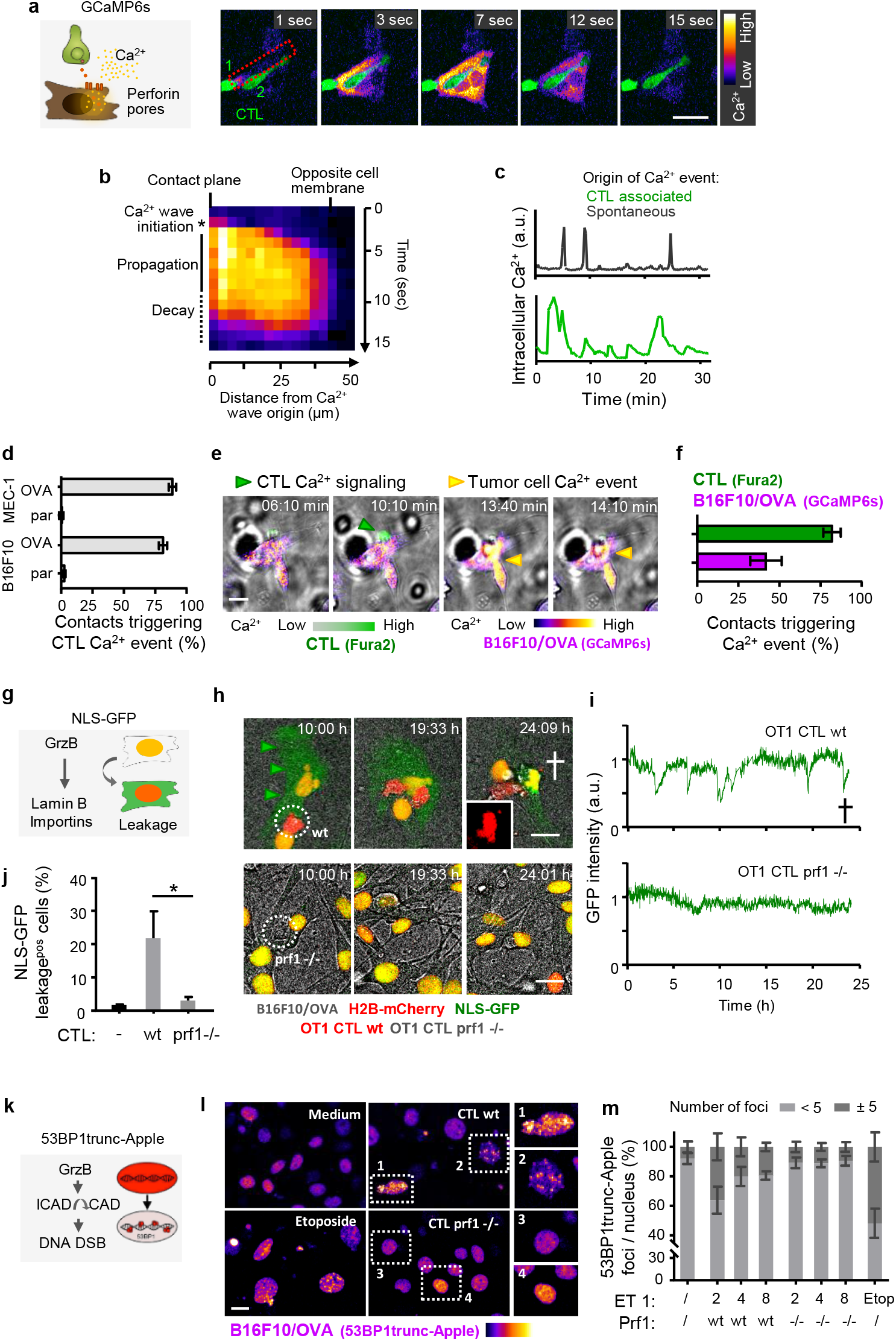
CTL-mediated sublethal damage kinetics. **a,** Principle of reporter assay (cartoon, left) and time-lapse sequence of OT1 CTL (green) associated Ca^2+^ event (Fire LUT) in MEC-1/OVA target cell. Frame rate: 1 sec. **b,** Ca^2+^ influx originating at the CTL-target cell contact plane and propagating through the target cell cytoplasm. **c,** Representative example of amplitude and duration of CTL-associated calcium events compared with spontaneous intracellular Ca^2+^ fluctuations in the absence of contacting CTL. **d,** Percentage of CTL contacts associated with Ca^2+^ events in OVA expressing target cells (OVA) compared to OVA negative parental (par) cells. Data show the means ± SD from 3 independent experiments. **e,** Image sequence of Fura2-labeled OT1 CTL during contact with GCaMP6s expressing B16F10/OVA cell. Arrowheads: green, Fura2-positive event; yellow, GCaMP6s-positive event. **f,** Percentage of contacts triggering Ca^2+^ events in OT-1 CTL or B16F10/OVA target cells, respectively. Quantification was performed by manual analysis from 78 contacts (4 independent experiments). **g,** Principle of the NLS-GFP reporter for monitoring sublethal damage. Abbreviations: GrzB, Granzyme B; Yellow nucleus resulting from overlapping emission of NLS-GFP and H2B-mCherry in the nucleus; red nucleus, persisting H2B signal after leakage of NLS-GFP into the cytoplasm. **h,** Representative image sequence of wt and prf1−/− OT1 CTL interacting with NLS-GFP expressing target cells. Dotted circle, CTL; white cross, target cell apoptosis; inset, condensed, apoptotic nucleus (H2B-mCherry). **i,** NLS-GFP signal intensity measured in the nucleus over time and outcome. The GFP signal was divided by the H2B-mCherry signal to normalize for focus drifts which affect signal intensity. Cross, nuclear fragmentation. **j,** Percentage of NLS-GFP leakage events in cultures without or with wt or prf1−/− OT1 CTL. Data show the means ± SD (3 independent experiments). **k,** Principle of the 53BP1trunc-Apple DNA damage reporter. Abbreviations: GrzB, Granzyme B; ICAD/CAD, Inhibitor of Caspase-activated DNase; DNA DSB, DNA double-strand breaks; **l,** Representative micrographs of 53BP1trunc-Apple expressing B16F01/OVA after incubation for 48 h with medium, Etoposide (25 μg/ml), or in coculture with wt or prf1−/− OT1 CTL at ET ratio of 1:4. Zoomed images, nuclei with or without 53BP1trunc-Apple foci. **m,** Percentage of 53BP1trunc-Apple expressing B16F10/OVA cells showing <5 or ≥5 53BP1trunc-Apple foci per nucleus after 48 h culture under different conditions. Foci per nucleus were counted using a custom ImageJ/FIJI script to segment nuclei based on Hoechst counterstaining as the number of maxima per nucleus (Find Maxima plugin). Data show the means ± SD (duplicates from 3 independent experiments). Scale bars, 20 μm.

**Extended data figure 4.**
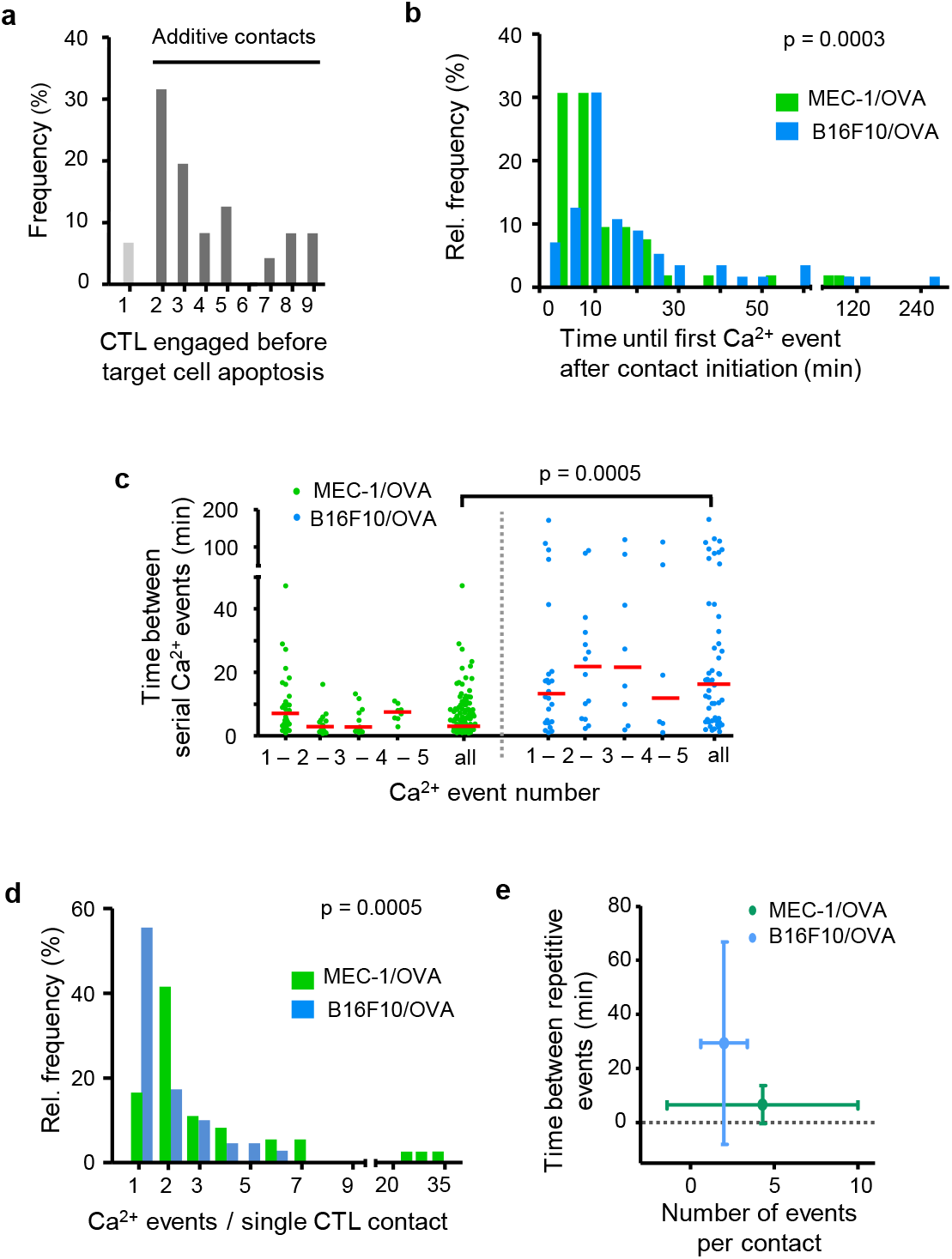
Kinetics and frequency of CTL-induced sublethal events. **a,** Percentage of CTL engaged before target cell death in B16F10/OVA coculture with OT1 CTL**. b,** Lag phase until first Ca^2+^ event after contact initiation in MEC-1/OVA and B16F10/OVA target cells. Data from 55 (B16F10) and 52 (MEC-1) Ca^2+^ events. p value, two-tailed Mann-Whitney test. **c,** Intervals between sequential Ca^2+^ events in the target cell induced by the same CTL in one single contact. Medians were 4 min (MEC-1/OVA) and 18 min (B16F10/OVA). Red line, median. Data show 53 (B16F10) and 119 (MEC-1) Ca^2+^ events. **d,** Number of Ca^2+^ events associated with the same CTL. Data show 55 (B16F10) and 36 (MEC-1) CTL. Data from (b-d) pooled from 3 (B16F10) and 2 (MEC-1) independent experiments. p values, two-tailed Mann-Whitney test. **e,** Number of Ca^2+^ events per CTL contact plotted against frequency of repetitive Ca^2+^ events in MEC-1/OVA cells compared to the B16F10/OVA cells. Data from 53 (B16F10) and 119 (MEC-1) Ca^2+^ events pooled from 3 (B16F10/OVA) and 2 (MEC-1/OVA) independent experiments.

**Extended data figure 5.**
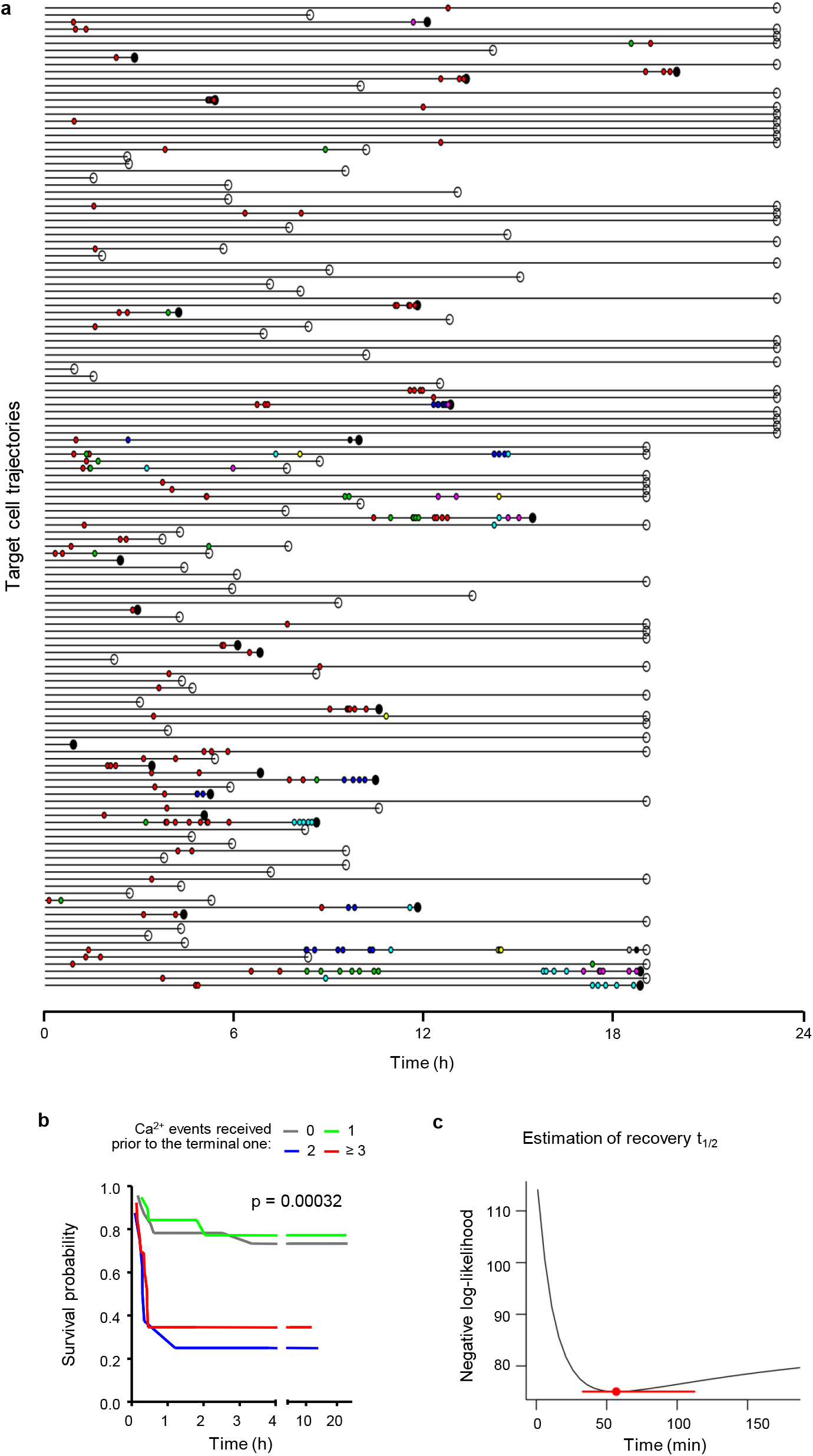
Statistical analysis. **a,** Time-points of Ca^2+^-positive events (dots color-coded for CTL causing the event) in individual B16F10/OVA cells contacted by OT-1 CTL. Outcome of the interaction is indicated as lethal (filled black dot) or non-lethal (open circle) at the end of each trajectory. **b**, Survival probability of B16F10/OVA cells having received increasing numbers of Ca^2+^ events prior to the terminal event after filtering redundant Ca^2+^ events. p-values, log rank test comparing all groups. **c,** Estimation of damage recovery half-life in B16F10/OVA after one single Ca^2+^ event by a statistical model that assumes additive killing (see Methods). Point and error bars: damage half-life that is most consistent with the data and 95% CI.

**Extended data figure 6.**
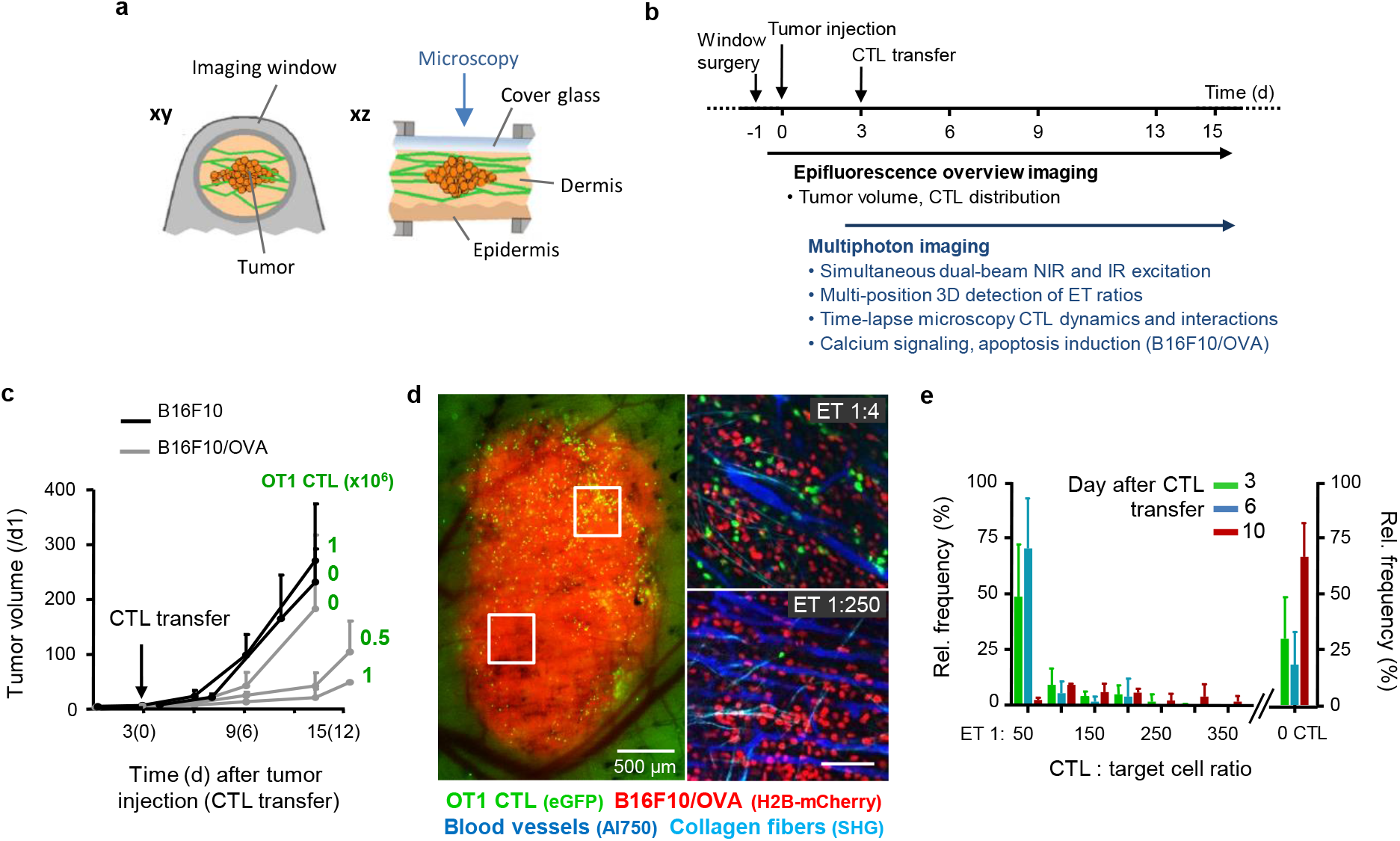
Intravital microscopy. **a,** Dorsal skin-fold chamber setup and tumor model. **b,** Setup and time line, including tumor cell injection, CTL i.v. transfer, intravital monitoring and parameter extraction. **c,** Time-dependent impact of adoptive OT1 CTL transfer on B16F10 tumor volume based on measurements obtained from epifluorescence overviews. Error bars, means ± SD (5 independent tumors). **d,** Tumor morphology and distribution of CTL monitored by epifluorescence detection through the skin window (left image) and multi-photon microscopy images (right images) recorded in different tumor subregions. Red, tumor nuclei (H2B/mCherry); green, OT1 CTL (eGFP); cyan, collagen fibers (SHG); blue, blood vessels (70kDa-dextran/Alexa750). **e,** Sub-regional quantification of ET ratios over time. Error bars, means ± SD (8 independent tumors).

**Extended data figure 7.**
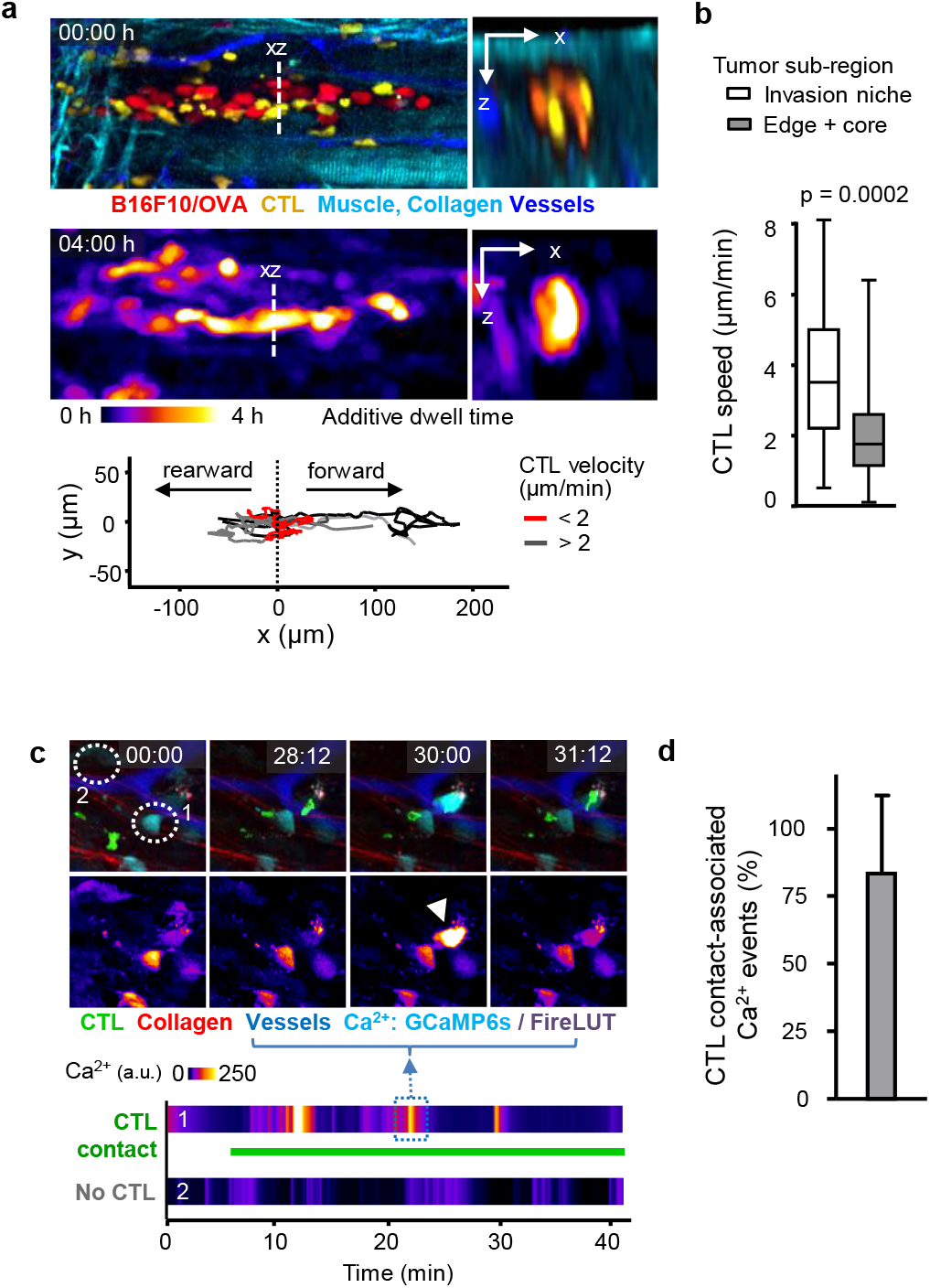
Monitoring dynamic cellular events by intravital time-lapse microscopy. **a,** Distribution (top), dwell-time (middle) and migration tracks (bottom) after 4 h time-lapse recording (top) of CTL (yellow) between tumor cells (red) along channel-like invasion niche confined by collagen bundles (cyan) and blood vessels (blue). Data from one representative time-lapse recording. **b,** CTL migration speed in relation to tumor subregions. Data shows 34 (invasion) and 120 (tumor edge) CTL tracks from 3 independent experiments. p value, two-tailed Mann-Whitney test. **c,** Sequential intracellular Ca^2+^ events (GCaMP6s) in B16F10/OVA cell during CTL engagement. Time stamp, min:sec. Green, OT1 CTL; Cyan, GCaMP6s in B16F10/OVA; Lower panel: Fire LUT, Ca^2+^ intensity (GCaMP6s) of either contact indicated by number. **d**, Percentage of Ca^2+^ events associated with a CTL contact. Error bars, mean ± SD, data from 47 Ca^2+^ events from 4 independent mice.

**Extended data figure 8.**
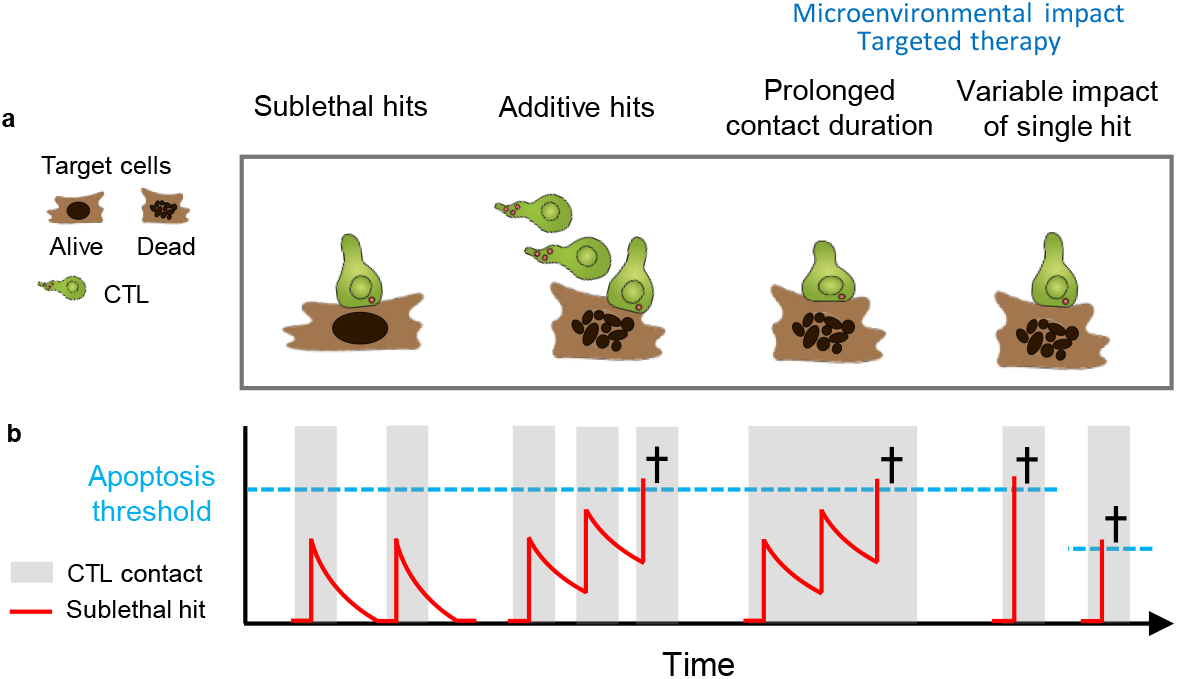
The concept of additive cytotoxicity. (a) CTL-target cell contact types and outcome. (b) Predicted accumulation of cytotoxic but sublethal hits and recovery over time. Black cross, target cell apoptosis.

