## Supplementary Method Section for "Cancer cell elimination by cytotoxic T cell cooperation and additive damage"

### Methods section

**Cell lines and primary cell culture.** C57BL/6 mouse embryonic fibroblast-derived cells (MEC-1) expressed B7.1 and the OVA-derived CTL epitope SIINFEKL (MEC-1/OVA) or the adenovirus type 5 E1A-derived CTL epitope SGPSNTPPEI (MEC-1/E1A) coupled to a signal-sequence which directs epitope expression to the endoplasmatic reticulum[1]. The cells were cultured in RPMI 1640 medium (GIBCO, 21875-034) supplemented with 10% FCS (SIGMA, F7524), 10 mM HEPES (GIBCO, 15630-056), 500 mM 2-mercaptopethanol, 1% penicillin and streptomycin (PAA, P11-010), 1% sodium pyruvate (GIBCO, 11360-039), and 0.1 mM non-essential amino acids (GIBCO, 11140-035).

B16F10 mouse melanoma cells expressing the OVA-derived CTL epitope SIINFEKL (B16F10/OVA) were transduced to express histone-2B/mCherry as described[2]. A low-pigmented subline was derived by sequential passaging and validated for unchanged growth, invasion ability, antigenicity and apoptosis resistance *in vitro* and *in vivo*. The cells were cultured in RPMI 1640 medium (GIBCO, 21875-034) supplemented with 10% FCS (SIGMA, F7524), 1% sodium pyruvate (GIBCO, 13360-039) and 1% penicillin and streptomycin (PAA, P11-010). Prior to experiments, cells were stimulated with 200 U/ml murine IFNγ (PEPROTECH, 315-05) for 48h.

MV3 and BLM male HLA-A2 expressing human melanoma cells were cultured in DMEM medium (GIBCO, 10938-025) supplemented with 10% FCS (SIGMA, F7524), glutamine (LONZA, BE17-605), 1% sodium pyruvate (GIBCO, 13360-039) and 1% penicillin and streptomycin (PAA, P11-010). Prior to experiments, cells were stimulated with 200 U/ml human IFNγ (SIGMA, I3265) for 48h.

The identity of B16F10, MCF-7, MV3 and BLM cells was verified by Short Tandem Repeat (STR) DNA profiling (IDEXX BioResearch). No respective mammalian or murine interspecies contamination was detected. Cells were routinely tested for mycoplasma contamination (MycoAlert, Lonza). All cell lines were negative for mycoplasma contamination.

**Mice.** C57BL/6 J mice (4-6 weeks of age) were purchased at Charles River Laboratories. Transgenic mice expressing eGFP under the human ubiquitin C promoter (Jackson Laboratories, C57BL/6-Tg(UBC-GFP)30Scha/J, stock number: 004353) and transgenic mice expressing dsRed under the chicken beta-actin promoter (Jackson Laboratories, STOCK Tg(CAG-DsRed*MST)1Nagy/J, stock number: 006051) were crossed to OT-1 TCR transgenic mice (Jackson Laboratories, C57BL/6-Tg(TcraTcrb)1100Mjb/J, stock number:

003831). Double transgenic eGFP/OT-1 and dsRed/OT-1 were bred in the Central Animal Laboratory of the Radboud University Nijmegen, The Netherlands.

C57BL/6-Prf1tm1Sdz/J (breeding pairs) were purchased at Jackson Laboratories (Stock No: 002407) and crossed to double-transgenic dsRed/OT-1 mice in the Central Animal Laboratory of the Radboud University Nijmegen, The Netherlands. Genotyping was performed by PCR following the genotyping protocol recommended by Jackson Laboratory and mice homozygous for mutant perforin-1 were used for experiments at 6 – 10 weeks of age.

**Isolation and activation of primary CD8$^+$ OT1 CTL.** OT1 CTL were activated as described[3]. In short, splenocytes from OT1 or double-transgenic eGFP/OT1 or dsRed/OT1 mice were isolated and erythrocytes were depleted by ammonium chloride (0.83% $NH_4Cl$, 0.1% $KHCO_3$, 0.37% $Na_2$ EDTA). For the expansion of antigen-specific CTL, splenocytes were cultured at a concentration of 2.5 x$10^5$ /ml in the presence of 0.5 µg/ml SIINFEKL peptide in 24-well plates for 3 days. On day 3, IL-2 (100 U/ml) (ABD SEROTEC, PMP-38) was added to the cultures. Lymphocytes were incubated for further 24-48 h and harvested on days 4-5 by a Ficoll gradient (AXIS-SHIELD PoC AS, Oslo, Norway). Purity was determined by flow cytometry and typically exceeded 96% of V$\alpha$2$^+$CD8$^+$CD62L$^{low}$CD44$^{hi}$ cells.

To determine the surface expression of Lamp-1 after 24h of coculture with target cells in the 3D cytotoxicity assay, CTL and surviving target cells were harvested by dissolving the collagen with collagenase I (40 U / 96-well; 30 min; SIGMA C0130) and detaching the remaining adherent cells with trypsin/EDTA (5 min). Both cell fractions were combined, washed in PBS, stained with AlexaFluor488-conjugated anti-Lamp-1 rat-IgG (BIOLEGEND, 121608) and detected after signal enhancement by donkey anti-rat/Alexa488 (LIFE TECHNOLOGIES, A21208). CTL were gated on intact morphology, viability by propidium iodide exclusion and dsRed expression using De Novo FCS Express 4. Percentages of positive events were calculated using the build-in FCS Express function for histogram subtraction.

**Activation and culture of primary human SMCY.A2 CTL.** The CD8$^+$ SMCY CTL line was isolated and cultured as described previously[2]. In short, CD8$^+$ CTL were isolated from PBMC obtained from a renal cell carcinoma patient who received allogeneic stem cell transplantation (SCT) and donor lymphocyte infusions (DLI). This patient has given informed consent to the prospective collection of peripheral blood samples for investigational use, which was approved by the Radboud University Medical Centre (RUMC) Institutional Review Board. CD8$^+$ T cells were expanded by weekly stimulation with PBMC obtained before SCT in Iscove's modified Dulbecco's medium (IMDM; INVITROGEN, Carlsbad,

CA) supplemented with 10% human serum (HS; Sanquin blood bank, Nijmegen, the Netherlands). After initial stimulation, CTL (0.5 x $10^6$) were cultured in IMDM/10% HS containing irradiated (80 Gy) recipient EBV-LCL (0.5 x $10^6$) and irradiated (60 Gy) allogeneic PBMC (0.5 x $10^6$) from two donors, together with IL-2 (100 IU/ml; Chiron, Emeryville, CA) and PHA-M (1 mg/ml; Boehringer, Alkmaar, the Netherlands). A greater than 95% pure population of SMCY.A2 specific CTL was isolated by FACS sorting (tetramer staining of HLA-A2-restricted SMCY.A2 epitope FIDSYICQV). Specificity of SMCY.A2 CTL was further confirmed by lack of cytotoxicity against female HLA-A2 expressing breast carcinoma cell (MCF-7) as target cells (Fig. 1h).

**Three-dimensional cytotoxicity assay and time-lapse microscopy.** A sub-confluent monolayer of target cells was overlaid with a 3D collagen gel (PureCol, concentration: 1.7 mg/ml) containing pre-activated CTL in a customized imaging chamber, allowed to polymerize (30 min, 37°C) and filled with undiluted CTL growth medium. CTL dynamics were recorded by time-lapse bright-field microscopy with a 30 sec frame interval for 24-48 h. CTL-target cell interactions were quantified by manual analysis. CTLs in contact with target cells slowed migration speed and spread on the target cell surface, which was used as morphological marker to distinguish irrelevant passenger CTL. For quantification of the frequency of serial killing events, only CTLs which could be followed for >12 h were included in the statistical analysis. CTL-target cell interactions were classified as non-lethal after survival of the target cell for > 3 h after CTL detachment. The > 3 h follow-up period is based on statistics of the lag time between last CTL induced $Ca^{2+}$ event and target cell death which predominantly remained below 1 h in MEC-1/OVA and 1-2 h in B16F10/OVA (Extended Data Fig. 5h).

**Monitoring sublethal damage in target cells.**

MEC1/OVA and B16F10/OVA target cells were lentivirally transduced to stably express the calcium sensor GCaMP6s[3], selected with blasticidin (10 µg/ml: Life Technologies, R210-01) and FACS-sorted on GCaMP6s background expression. pCDH-NLS-copGFP-EF1-BlastiS coding for the NLS-GFP[4] reporter was a gift from Jan Lammerding (Addgene plasmid # 132772; http://n2t.net/addgene:132772; RRID:Addgene_132772). Histone-2B-mCherry expressing B16F10/OVA cells were lentivirally transduced to co-express NLS-GFP, selected for stable expression with blasticidin (10 µg/ml: Life Technologies, R210-01) and FACS-sorted for double-positive GFP and mCherry expression. Apple-53BP1trunc[5] was a gift from Ralph Weissleder (Addgene plasmid # 69531; http://n2t.net/addgene:69531;RRID: Addgene_69531). B16F10/OVA cells were lentivirally transduced to stably express Apple-

53BP1trunc, selected for stable construct integration with puromycin (2 µg/ml: Sigma-Aldrich, P7255) and FACS sorted for Apple positive cells. Cells were further subcloned and 3 clones were pooled to achieve uniform reporter expression levels.

*Time-lapse monitoring of sublethal damage*. CTL-target cell conjugation and associated sublethal damage were monitored by co-registering GCaMP6 and dsRed OT1 CTL at frame intervals of 8-12 sec for up to 12 h (GCaMP6s) or frame rates of 2 min for up to 30 h (NLS-GFP). For monitoring CTL conjugation with Apple-53BP1trunc-expressing target cells which spectrally overlaps with dsRed2, transmission contrast of unlabeled wt OT1 CTL and fluorescence of Apple-53BP1trunc were recorded at time intervals of 5 - 10 min for up to 48 h (Leica SP8 SMD Cconfocal). Excitation was limited to 0.05 mW for each excitation line (561 nm for Histone-2B-mCherry, Apple-53BP1trunc and dsRed2; 488 nm for GCaMP6s and NLS-GFP). Viability of CTL and target cells during long-term imaging was controlled by verifying constant CTL migration speed and morphological integrity over time, lack of laser induced cell death, and unperturbed proliferation compared to bright-field imaging in cell culture without CTL. CTL-associated damage events were identified by manual or semi-automated image segmentation and intensity analysis using ImageJ/FIJI.

$Ca^{2+}$ signals in OT1 CTL and target cells were monitored by spinning-disk confocal microscopy (BD Pathway) during 3D coculture of Fura2-labeled OT1 CTL with GCaMP6s expressing B16F10/OVA cells, using frame rates of 95 sec for Fura2 (340 / 380 nm) and 8 sec for GCaMP6s excitation (488 nm). To account for phototoxicity and bleaching, imaging periods were limited to 1 h.

**Intravital multiphoton microscopy.** All experiments were approved by the Ethical Committee on Animal Experiments and performed in the Central Animal Laboratory of the Radboud University, Nijmegen (RU-DEC 2009-174, 2011-298, 2017-0034), in accordance with the Dutch Animal Experimentation Act and the European FELASA protocol (www.felasa.eu/guidelines.php).

Histone-2B/mCherry expressing tumor cells ($1x10^5$) were injected into the deep dermis of C57/B16 J mice (Charles River) carrying a dorsal skin-fold chamber and were repeatedly monitored for up to 15 days[6]. Three days after tumor implantation, *in vitro* activated dsRed OT1 CTLs ($0.5–1x10^6$) were injected intravenously. Multi-parameter intravital multiphoton microscopy was performed on anesthesized mice (1-3% isoflurane in oxygen) on a temperature-controlled stage (37°C). Blood vessels were contrasted by intravenous injection of AlexaFluor750-labeled 70kD dextran (2 mg/mouse; Invitrogen).

Tumor volume was reconstructed from epifluorescence overview images recorded at sequential time points. Tumor volume (V) was calculated as (tumor width)$^2$ x (tumor length) x $\pi/6$.

Imaging was performed on a customized near-infrared/infrared multiphoton microscope (TriMScope-II, LaVision BioTec, Bielefeld, Germany), equipped with three tunable Ti:Sa (Coherent Ultra II Titanium:Sapphire) lasers and an Optical Parametric Oscillator (OPO). 4D time-lapse recordings of CTL interactions with tumor cells were acquired by sequential scanning with 910 nm (GFP, Al750) and 1090 nm (mCherry, dsRed2, SHG) at a laser power of 30 mW (910 nm) and 60 mW (1090 nm) with a sampling rate of 1 frame / 2 min over periods of 4-8 h. For visualization of intracellular $Ca^{2+}$ dynamics, 3D volumes of 250 x 250 x 100 µm were acquired by simultaneous excitation at 910 nm (GCaMP6s, Al750; 20 mW) and 1140 nm (mCherry, dsRed2, SHG; 30 mW) with a sampling rate of 1 frame / 10-15 sec over periods of 1-2 h.

**Image processing and quantification.** Images were processed using Fiji/ImageJ[7] (http://pacific.mpi-cbg.de/wiki/index.php/Fiji). Mosaic images were stitched using the Stitch Grid/Collection plugin[8] and drifts in time-lapse recordings were corrected using the StackReg plugin[9] and the Correct 3D Drift plugin. For presentation of time-lapse recordings, 4D stacks were adjusted for bleaching using the Bleach Correction plugin (Histogram normalization). If necessary background noise was filtered using the Remove Outliners plugin and images were scaled and adjusted for brightness, contrast and gamma to enhance visualization. For image quantifications the raw, unmodified images were used.

Intact vs. apoptotic states of tumor nuclei and CTL ratios were quantified from individual slices of multiple 3D stacks of 350 x 350 µm, acquired with 7 µm z-resolution until imaging depths of up to 300 µm. To avoid repeated counting of the same cell, every 3$^{rd}$ slice per stack was analyzed. Nuclei and CTL were segmented by applying a Gaussian Blur filtering followed by automated thresholding (Li algorithm) and separation of touching objects using Watershed. Using the Analyze Particles command of FIJI, segmented objects were filtered for size (Nuclei: $\geq$ 60 µm$^2$; CTL: $\geq$ 50 µm$^2$) and counted per slice. To determine subregional apoptosis and mitosis rates, apoptotic and mitotic nuclei were counted manually and calculated as percentage of total nuclei per slice.

53BP1trunc-Apple foci were analyzed using custom-scripts to segment nuclei based on Hoechst counterstaining, followed by the detection and counting of foci per nucleus in the Apple Channel using the Find Maxima plugin in FIJI/ImageJ.

NLS-GFP leakage events were quantified by manual tracking of the tumor nuclei using the Manual Tracking Plugin. The intensity of nuclear GFP was divided by the H2B-mCherry signal to correct for mild focus drifts which affect GFP intensity. Normalized GFP values per nucleus were plotted over time and analyzed manually for leakage events, in combination with manual inspection of the time-lapse recording.

**Statistical modeling.** The analysis of target cell survival, $Ca^{2+}$ events and the resulting survival probability curves were computed in GNU R using the 'survival' and 'rms' packages. In a subset of analyses, redundant $Ca^{2+}$-positive “hits” which may bias the analysis by concealing the already occurred lethal hit and being misleadingly interpreted as prior, additive hits were removed, based on the following considerations. The "stochastic killing" hypothesis predicts that each $Ca^{2+}$ event has the same likelihood to induce apoptosis and, consequently, the number of $Ca^{2+}$ events received prior to the last event preceding apoptosis should not impact target cell survival. However, redundant $Ca^{2+}$ events induced by the same or other CTL, which are dispensable for apoptosis, may occur after target cells received the lethal hit, and result on an overestimation of required hits. We used the following approach to identify and remove such redundant events. Waiting times to apoptosis following the last lethal event were assumed to be exponentially distributed, and the mean waiting time best describing the data was fitted using the maximum likelihood method. For each target cell, the $Ca^{2+}$ event whose likelihood of being the terminal event was largest, given the fitted distribution, was determined. This event was defined as the terminal event for the given cell, and all subsequent $Ca^{2+}$ events were considered redundant (Extended Data Fig. 5a). Despite the removal of redundant contacts, survival probability and lag times to apoptosis remained significantly dependent on prior $Ca^{2+}$ events ($p=0.00032$ with removal, $p=0.00003$ without), in agreement with additive cytotoxicity of serial $Ca^{2+}$ events (Extended Data Fig. 5b).

To simulate purely stochastic killing, the time between serial $Ca^{2+}$ events and the lag time between $Ca^{2+}$ events to apoptosis were randomly permuted. Assuming stochastic killing, these permutations should not affect target cell survival probability after the last contact. However, the permuted data revealed significantly enhanced killing efficiency compared to the actual killing efficiency derived from live-cell measurements whereby the amount of prior $Ca^{2+}$ events lacked impact on survival probability after the last $Ca^{2+}$ event (Fig. 3d). Thus, stochastic apoptosis induction by individual hits is incompatible with the experimentally observed killing kinetics.

Cell damage repair times were estimated from the $Ca^{2+}$ imaging data in target cells using a Cox proportional hazards model based on the following assumptions: (1) Each $Ca^{2+}$ event induces a unit amount of damage to the cell; (2) existing damage decays exponentially; (3) the instantaneous risk of apoptosis (hazard) is proportional to the current amount of damage. The damage was entered into the model as a time-dependent covariate and its coefficient represents the amount of damage dealt by each hit. The decay rate of the exponential function was then estimated by minimizing the negative log-likelihood of the model.

**Statistical analysis.** Unpaired student's t-tests or Mann-Whitney U-tests, as appropriate, were applied using GraphPad Prism 5.

#### References Material and Methods
